# Cell-type specific sensory and motor activity in the cuneiform nucleus and pedunculopontine nucleus in mice

**DOI:** 10.1101/2025.03.25.645329

**Authors:** Cornelis Immanuel van der Zouwen, Andrea Juárez Tello, Jacinthlyn Sylvia Suresh, Juan Duque-Yate, Ted Hsu, Vaibhav Konanur, Joël Boutin, Mitchell F. Roitman, Dimitri Ryczko

## Abstract

The activity of neurotransmitter-based cell types in the cuneiform and pedunculopontine nuclei during locomotion, non-locomotor behaviors, and following sensory stimulation is not fully understood. Using fiber photometry in mice, we found cell-type specific responses to sensory stimuli. Glutamatergic and GABAergic cells responded to sound, visual looming, and air puffs, except for pedunculopontine GABAergic cells, which did not respond to visual looming. Cholinergic cells responded to air puffs. When a stimulus triggered high-speed locomotion, activity increased in cuneiform glutamatergic neurons. Conversely, when low-speed locomotion was triggered, activity increased in pedunculopontine glutamatergic neurons. During spontaneous low-speed locomotion, activity increased in pedunculopontine glutamatergic cells. Activity also increased in a cell type-specific manner during grooming or rearing. Our study shows cell type-specific activity in the cuneiform or pedunculopontine nuclei during locomotion, non-locomotor behaviors, and following sensory stimulation. Sensory responsiveness likely has relevance in Parkinson’s disease, where sensory circuits are increasingly targeted to improve walking.

## INTRODUCTION

The Mesencephalic Locomotor Region (**MLR**) plays a key role in the control of locomotion from lamprey to mammal (for review^1–4^). In mammals, the anatomical nuclei corresponding to the MLR include the cuneiform nucleus (**CnF**) and pedunculopontine nucleus (**PPN**). These nuclei contain altogether glutamatergic, GABAergic and cholinergic neurons. Studies based on optogenetics and chemogenetics established that neurotransmitter-based cell types in the CnF and PPN control different aspects of locomotion. CnF glutamatergic neurons (vesicular glutamatergic transporter 2-positive, **Vglut2^+^**) initiate locomotion and control speed^5–9^. CnF GABAergic neurons (vesicular GABA transporter-positive, **VGAT^+^**) stop locomotion^5^. The role of PPN neurons is more complex, probably because of their intricate projection pattern (for review^10,4^). PPN Vglut2^+^ neurons have been shown to either initiate locomotion^5,11^ or to stop locomotion^6,7^. PPN VGAT^+^ neurons were reported to either stop locomotion^5,12^ or to evoke locomotion flanked with numerous stops^11^. PPN cholinergic neurons (choline acetyltransferase-positive, **ChAT^+^**) have been found to increase, decrease, or to have no effect on locomotion^13,14,5,6,7,15^ (for review^4^).

The activity of genetically defined-cell types in the CnF or PPN during locomotion is not yet fully understood. Extracellular recordings of CnF and PPN Vglut2^+^ neurons in head-fixed mice walking on a motorized treadmill showed a positive correlation between speed and firing frequency, with PPN neurons showing greater selectivity at low speed^5^. Similar observations were reported for some Vglut2^+^ or calcium/calmodulin-dependent protein kinase IIa promoter-positive (**CamKIIa^+^**) neurons recorded without distinction in the CnF or PPN in head-fixed mice walking on a trackball^16,13^. GABAergic neurons recorded without distinction in the CnF or PPN can encode either a locomotor or stationary state, highlighting the complexity of this population^13^. Electrophysiological recordings also show that unidentified PPN neurons correlate positively or negatively with speed in rats walking in an open field^17^ or an elevated maze^18^. In zebrafish, neurons in a region corresponding to the MLR exhibit increased calcium signals during swimming^19^. Beyond locomotion, some glutamatergic cells in the CnF, PPN or neighboring mesencephalic reticular formation are active during handling, grooming or rearing in mice^20^. However, a complete overview of Vglut2^+^, VGAT^+^ or ChAT^+^ neuron activities in the CnF or PPN during self-paced locomotion and other motor behaviors is still lacking.

The activity of CnF and PPN cells in response to sensory inputs is also not completely understood. Although the MLR is traditionally considered as a motor structure, locomotion potentially needs to be initiated in various environmental contexts, which could be channeled to the MLR through sensory modalities. In lamprey for instance, olfactory information can evoke locomotion through a synaptic relay in the MLR^21,22^. In mammals, the CnF is part of a system regulating defensive behaviors such as fleeing or freezing in response to a threat (for review^23,2^). The PPN is part of an exploratory system at the interface between the basal ganglia and the motor circuits (for review^23,10,1^). However, it is not fully understood whether different sensory stimuli evoke cell type-specific responses in the CnF or PPN, and whether activation of a specific cell type in these nuclei is associated with sensory-evoked locomotion.

Here, using fiber photometry in freely moving mice, we recorded Vglut2^+^, VGAT^+^ and ChAT^+^ neurons in the CnF and PPN during motor behaviors (self-paced spontaneous locomotion, grooming, rearing) and in response to sensory stimulations (sound, visual looming, or air puff). During spontaneous low-speed locomotor bouts, PPN Vglut2^+^ activity increased. When high-speed locomotion was triggered by air puffs, activity was increased in CnF Vglut2^+^ neurons. Conversely, when low-speed locomotion was triggered by sound and visual looming, activity was increased in PPN Vglut2^+^ neurons. During grooming and rearing, cell-type specific patterns of activity were found. Furthermore, each cell type displayed a specific response pattern to sound, visual looming or air puff, with CnF and PPN Vglut2^+^ neurons, and CnF VGAT^+^ neurons, being the most responsive. The sensory responsiveness of those cells likely has clinical relevance because sensory circuits are increasingly targeted to improve locomotor function in Parkinson’s disease (for review^24,25^).

## RESULTS

### Sound-evoked Ca^2+^ activity in CnF and PPN neurons

To record *in vivo* the Ca^2+^ activity of genetically-defined cell types, we injected an adeno-associated virus (**AAV**) driving the expression in a cre-dependent manner of the calcium sensor jGCaMP7f^26^ in the CnF or PPN of Vglut2-cre^27^, VGAT-cre^27^ or ChAT-cre mice^28^. We implanted an optic fiber ∼150 µm above the injection site and recorded Ca^2+^ activity in freely moving mice using fiber photometry^29^ (Figure S1). We exposed the mice to a clinking metallic sound (86.9 ± 0.5 dB, sound spectrogram illustrated in Figure S2A,S2B, see Methods) generated ∼1 m away from the mouse moving freely in an open-field arena (Figure 1A). Such sound increased the mean Ca^2+^ signal during the 5 s after sensory stimulation compared to the 5 s before in CnF Vglut2^+^ cells (Figures 1B,1G,1H,1M,1N, n = 7 mice), in CnF VGAT^+^ cells (Figures 1C,1I,1O, n = 7 mice), in PPN Vglut2^+^ cells (Figures 1D,1J,1P n = 9 mice), in PPN VGAT^+^ cells (Figures 1E,1K,1Q, n = 10 mice), but not in PPN ChAT^+^ cells (Figures 1F,1L,1R, n = 9 mice). Overall, this indicates that glutamatergic and GABAergic neurons in the CnF and PPN responded to sound, but not PPN cholinergic neurons.

**Figure 1.**
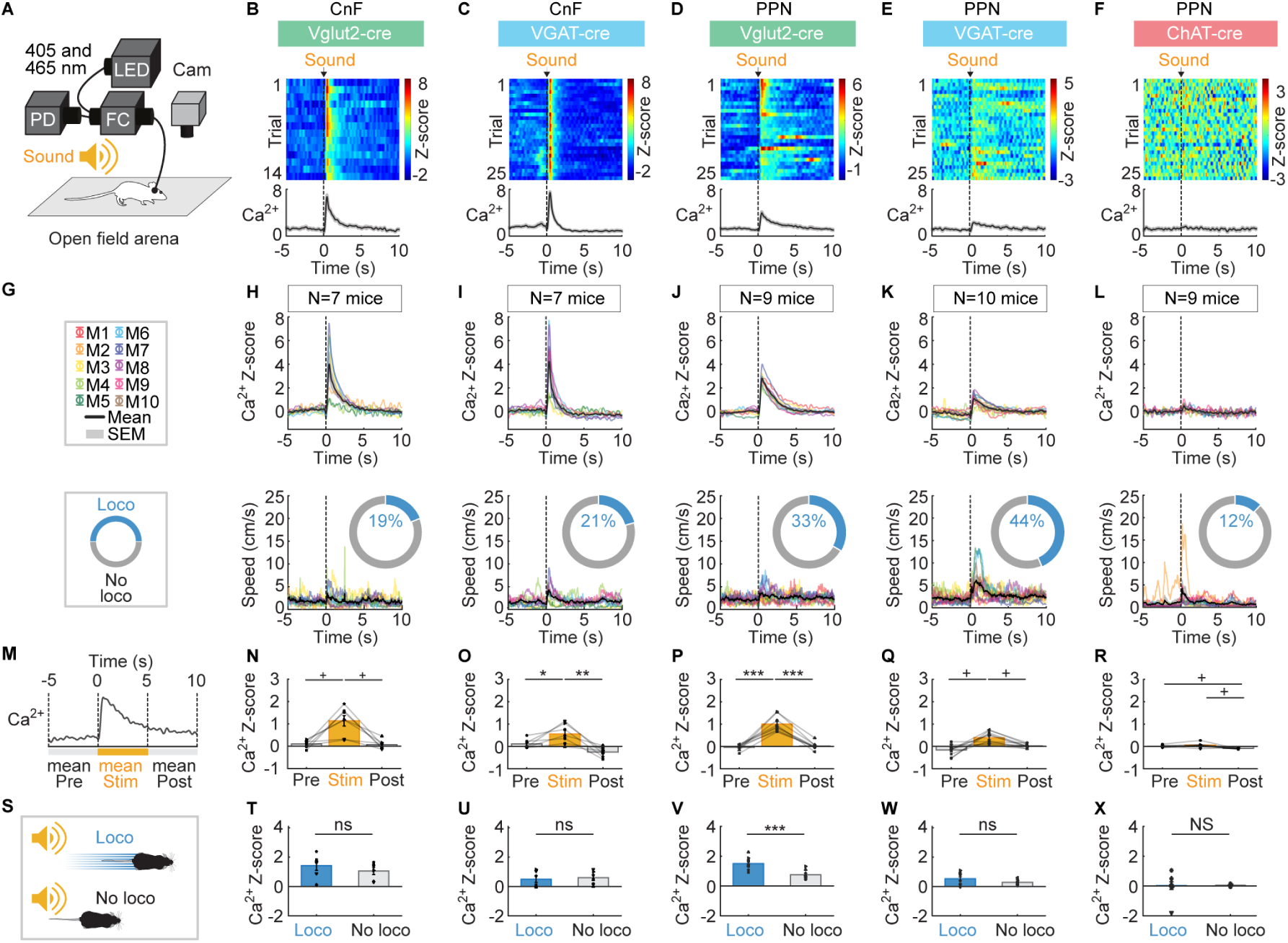
Sound-evoked responses in CnF and PPN neurons. (A) Vglut2-cre, VGAT-cre or ChAT-cre mice were injected in the cuneiform nucleus (**CnF**) or pedunculopontine nucleus (**PPN**) with an adeno-associated virus encoding for the genetically encoded calcium (**Ca^2+^**) indicator jGCaMP7f in a cre-dependent manner (see Methods) and implanted with an optic fiber ∼150 μm above the injection site (Figure S1). Fiber photometry recordings were done using two LEDs for isosbestic and jGCaMP7f signals (405 and 465 nm), a filter cube (**FC**) and a photodetector (**PD**). A clinking metallic sound (86.9 ± 0.5 dB, spectrogram in Figures S2A-S2C) was applied every 60.1 s ± 0.3 s, ∼1 m away from an open-field arena (see Methods). (B-F) Top, Ca^2+^ responses in the CnF or PPN evoked by sound stimulation. Each line illustrates a trial. Sound stimulus onset is indicated by a black vertical dotted line. Ca^2+^ responses are expressed in Z-score (see Methods), with warmer colors indicating stronger responses. Bottom, mean Z-scores ± SEM are illustrated. (G-L) Sound-evoked mean Ca^2+^ response ± SEM (top) and locomotor speed ± SEM (bottom) obtained in each mouse (M) (1-4 series of 4-10 trials per animal). The number of mice (N) used is indicated. The colored rings illustrate the mean proportion of trials in which mice displayed a locomotor response (speed > 3 cm/s for at least 0.5 s^8,9^) during the 5 s after sound stimulation. (M-R) mean Ca^2+^ signal before (-5 to 0 s), during (0 to 5 s) and after sound stimulation (5 to 10 s) (9-36 trials per animal; \**P* < 0.05, \*\**P* < 0.01, \*\*\**P* < 0.001, Student-Newman-Keuls test after a significant one-way repeated measures ANOVA; ^+^*P* < 0.05, Student-Newman-Keuls test after a significant Friedman ANOVA on ranks). (S-X) Comparison of mean Ca^2+^ signals in trials with (blue) or without (grey) a locomotor response during the 5 s following sound stimulation (ns, not significant, \*\*\**P* < 0.001 paired t-test; NS, not significant, Wilcoxon test; mice without locomotor trials or without non-locomotor trials were excluded from the analysis).

Increases in locomotor activity (speed > 3 cm/s for more than 0.5 s)^8,9^ were sometimes evoked during the 5 s after onset of the clinking metallic sound (12-44% of stimulations, Figures 1G-2L). Sound-evoked mean Ca^2+^ signals were significantly higher in PPN Vglut2^+^ cells if a locomotor response was induced (Figure 1S,1V). Such Ca^2+^ signal increase was not observed in the other CnF and PPN cell types during those trials where sound induced locomotion (Figures 1S-1U,1W-1X). Interestingly, Ca^2+^ responses were evoked in trials without detectable locomotor activity occurring at that time (including e.g. in CnF Vglut2^+^ neurons that are known to drive locomotion when activated optogenetically^5–8^).

**Figure 2.**
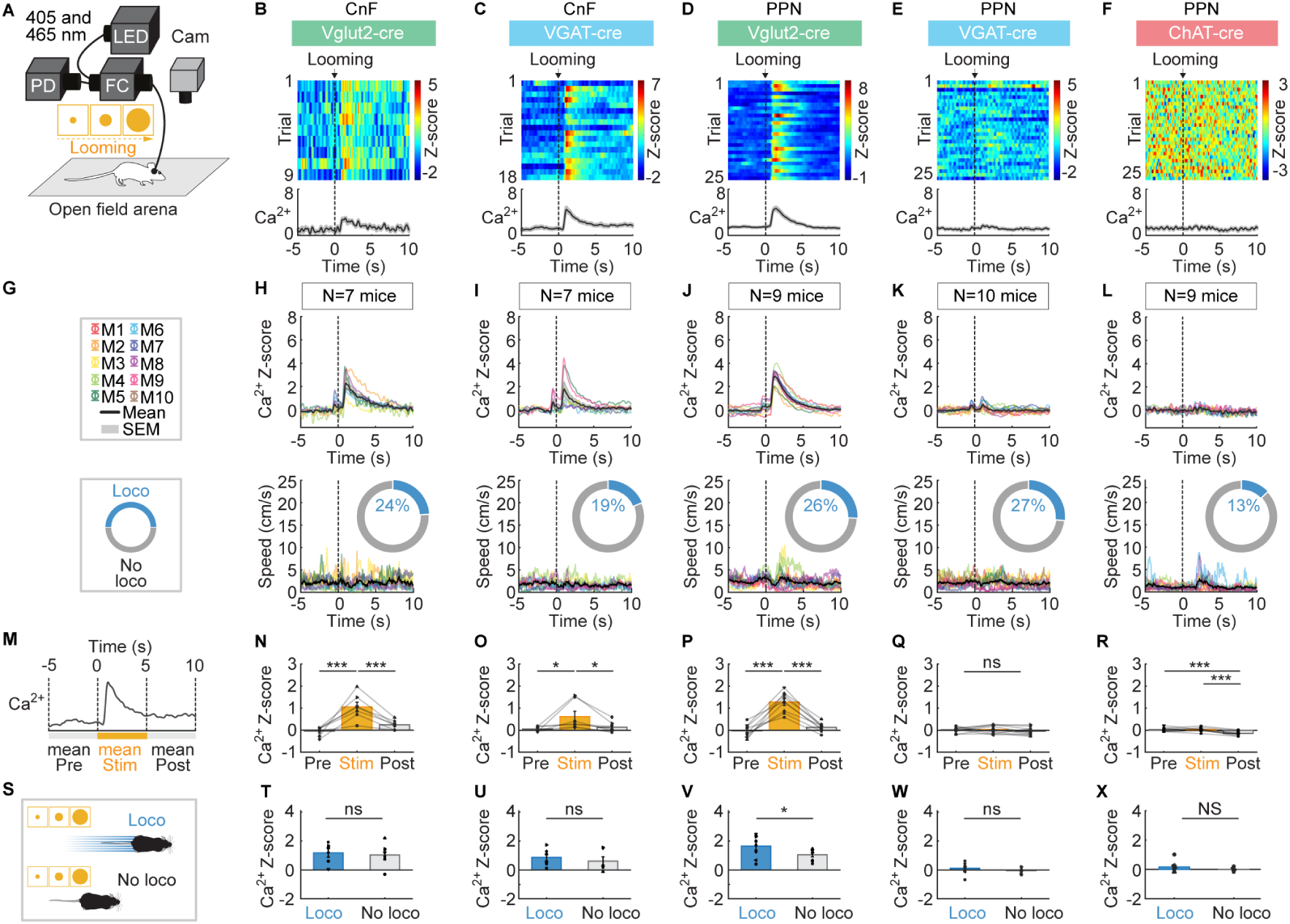
Visual-looming-evoked responses in CnF or PPN neurons. (A) Fiber photometry recordings were done as in Figure 1. Visual looming stimulus (a black disk increasing in diameter on a grey background during 1.676 s, see disk expansion rates provided in Figure S3) (see Methods) were applied every 61.1 s ± 0.5 s on a screen placed laterally relative to an open-field arena. (B-F) Calcium (**Ca^2+^**) responses evoked by visual looming stimuli in the CnF or PPN cells for example animals color-coded as in Figure 1. Each line illustrates a trial. The black vertical dotted line corresponds to t = 0.803 s after the beginning of the looming stimulus, when disk expansion starts to increase exponentially (see Figure S3). Bottom, mean Z-scores ± SEM are illustrated. (G-L) Visual looming-evoked mean Ca^2+^ response ± SEM (top) and locomotor speed (bottom) obtained in each mouse (M) (1-3 series of 8-10 trials per animal). The number of mice (N) used is indicated. The colored rings illustrate the mean proportion of trials in which mice displayed a locomotor response defined as in Figure 1 (see Methods). As for B-F, the black vertical dotted lines correspond to the start of the looming stimulus, which occurs 0.803 s after the start of the video with the stimulus. Note that the small increase in photometry signal before the black vertical dotted line was caused by our synchronization procedure, which relied on a button press that generated a small noise (see Figure S2C-S2F) (see Methods). (M-R) Mean Ca^2+^ signal before (-5 to 0 s), during (0 to 5 s) and after visual looming stimulation (5 to 10 s) (8-28 trials per animal; ns, not significant, \**P* < 0.05, \*\*\**P* < 0.001, Student Newman-Keuls test after a significant one-way repeated measures ANOVA). (S-X) Comparison of Ca^2+^ signals in trials with or without a locomotor response during the 5 s following visual looming stimulation (ns, not significant, \**P* < 0.05 paired t-test; NS, not significant, Wilcoxon test; mice without locomotor trials or without non-locomotor trials were excluded from the analysis).

### Visual looming-evoked Ca^2+^ activity in CnF and PPN neurons

We exposed the mice in the open-field arena to a visual looming stimulus, i.e. a black disk increasing in diameter during ∼1.7 s over a grey screen placed on the side of the open-field arena (Figure 2A; disk expansion rates illustrated in Figure S3A-S3C, see Methods). Visual looming increased the mean Ca^2+^ signal during the 5 s after sensory stimulation compared to the 5 s before in CnF Vglut2^+^ cells (Figures 2B,2G,2H,2M,2N, n = 7 mice), in CnF VGAT^+^ cells (Figures 2C,2I,2O, n = 7 mice) and in PPN Vglut2^+^ cells (Figures 2D,2J,2P, n = 9 mice), but did not change Ca^2+^ signal in PPN VGAT^+^ cells (Figures 2E,2K,2Q, n = 10 mice) or in PPN ChAT^+^ cells (Figures 2F,2L,2R, n = 9 mice).

Increases in locomotor activity were sometimes evoked during the 5 s after onset of visual looming (13-27% of stimulations, Figures 2G-2L). As for sound stimulations, visual looming-evoked Ca^2+^ signals were significantly higher in PPN Vglut2^+^ cells if a locomotor response was induced (Figures 2S,2V). Such Ca^2+^ signal increase was not observed in the other CnF and PPN cell types during those trials where visual looming induced locomotion (Figures 2S-2U,2W-2X). As for sound stimulations, visual looming stimuli could evoke Ca^2+^ responses in trials without locomotor activity occurring at that time.

### Air puff-evoked Ca^2+^ activity in CnF and PPN neurons

We exposed freely moving mice to air puffs in an open-field arena (Figure 3). In addition to the somatosensory component of the air puff delivered on the back of the animal, the puff also generated a sound (55.2 ± 0.5 dB, sound spectrogram illustrated in Figure S2B,S2E, see Methods), and involved the visual component of a human hand holding the air bulb, making the air puff a multisensory stimulus. Air puffs significantly increased the mean Ca^2+^ signal during the 5 s after sensory stimulation compared to the 5 s before in CnF Vglut2^+^ cells (Figures 3B,3G,3H,3M,3N, n = 8 mice), in CnF VGAT^+^ cells (Figures 3C,3I,3O, n = 9 mice), in PPN Vglut2^+^ cells (Figures 3D,3J,3P, n = 9 mice), in PPN VGAT^+^ cells (Figures 3E,3K,3Q, n = 10 mice), and in PPN ChAT^+^ cells (Figures 3F,3L,3R, n = 9 mice).

**Figure 3.**
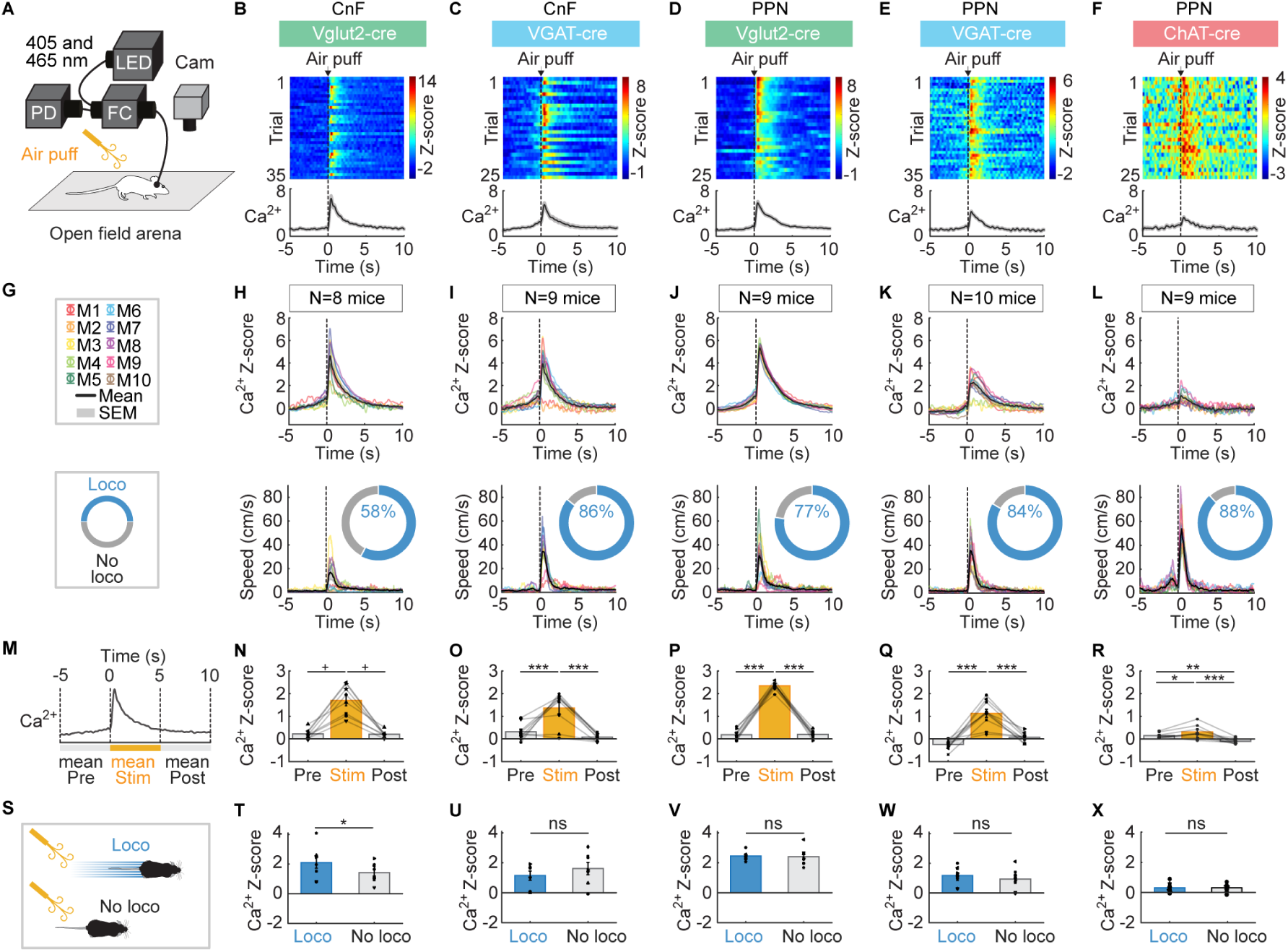
Air-puff-evoked responses in CnF and PPN neurons. (A) Fiber photometry recordings were done as in Figure 1. Air puff was applied with a small air bulb on the back of the animal every 61.4 s ± 0.7 s (see Methods). (B-F) Calcium (Ca^2+^) responses evoked by air puffs in the CnF or PPN cells for example animals color-coded as in Figure 1. Each line illustrates a trial. Air puff onset is indicated by a black vertical dotted line. Bottom, mean Z-scores ± SEM are illustrated. (G-L) Air puff-evoked mean Ca^2+^ response ± SEM (top) and locomotor speed ± SEM (bottom) obtained in each mouse (M) (2-4 series of 6-10 trials per animal). The number of mice (N) used is indicated. The colored rings illustrate the mean proportion of trials in which mice displayed a locomotor response defined as in Figure 1 (See Methods). (M-R) Mean Ca^2+^ signal before (-5 to 0 s), during (0 to 5 s) and after the air puff (5 to 10 s) (16-36 trials per animal; \**P* < 0.05, \*\**P* < 0.01, \*\*\**P* < 0.001, Student-Newman-Keuls test after a significant one-way repeated measures ANOVA; ^+^*P* < 0.05, Student-Newman-Keuls test after a significant Friedman ANOVA on ranks). (S-X) Comparison of mean Ca^2+^ signals in trials with or without a locomotor response during the 5 s following air puff (ns, not significant, \**P* < 0.05, paired t-test; mice without locomotor trials or without non-locomotor trials were excluded from the analysis).

Increases in locomotor activity were frequently evoked during the 5 s after air puff onset (58-88% of stimulations, Figures 3G-3L). In CnF Vglut2^+^ cells, Ca^2+^ signals were significantly higher if a locomotor response was evoked compared to trials without locomotor response (Figure 3S,3T). Such Ca^2+^ increase was not observed in the other CnF and PPN cell types during those trials where air puffs induced locomotion (Figures 3S,3U-3X). As for sound and visual looming stimulations, Ca^2+^ responses could be evoked by air puffs in trials without locomotor activity occurring at that time.

Interestingly, we found significant differences in mouse speed during locomotor bouts evoked by different sensory stimuli (*P* < 0.001, one-way repeated measures ANOVA, n = 37 mice with locomotor bouts evoked by all 3 stimuli pooled from all cell types). Speed was higher during air puff-evoked locomotor bouts (32.8 ± 1.3 cm/s) than during sound-evoked locomotor bouts (15.8 ± 0.7 cm/s; *P* < 0.001, Student-Newman-Keuls test) or visual looming-evoked locomotor bouts (16.3 ± 1.0 cm/s, *P* < 0.001, Student-Newman-Keuls test). Altogether, these findings suggest that CnF Vglut2^+^ cells are more active during the faster locomotor responses evoked by air puffs, while PPN Vglut2^+^ cells are more active during the slower locomotor responses evoked by sound or visual looming stimuli.

### Habituation to sensory stimulations

We observed habituation to sensory stimulations in some cell types (Figure S4). For sound, we found a negative linear relationship between stimulation trial number and mean Ca^2+^ signal in CnF Vglut2^+^ cells (*P* < 0.05, R^2^ = 0.64, -47% decrease at 9^th^ trial compared to 1^st^ trial, Figure S4B) and in PPN Vglut2^+^ cells (*P* < 0.05, R^2^ = 0.50, -67% decrease at 9^th^ trial, Figure S4D). For visual looming, we found a negative linear relationship between stimulation trial number and mean Ca^2+^ signal in CnF VGAT^+^ cells (*P* < 0.05, R^2^ = 0.58, -60% decrease at 9^th^ trial, Figure S4I). For air puff, we found a negative linear relationship between stimulation trial number and mean Ca^2+^ signal in CnF Vglut2^+^ cells (*P* < 0.01, R^2^ = 0.77, -60% decrease at 9^th^ trial, Figure S4N), in PPN Vglut2^+^ cells (*P* < 0.05, R^2^ = 0.62, -33% decrease at 9^th^ trial, Figure S4P), and in PPN VGAT^+^ cells (*P* < 0.05, R^2^ = 0.48, -31% decrease at 9^th^ trial, Figure S4Q). Such decreases of responsiveness over time likely reflects habituation to stimuli that did not represent any threat for survival.

### Ca^2+^ activity in CnF and PPN neurons during spontaneous, self-paced locomotion

We recorded the Ca^2+^ signals in the CnF and PPN cell types during bouts of spontaneous locomotion in an open-field arena (Figure 4). Locomotor bouts lasted 1.1 ± 0.02 s (at a speed of 11.9 ± 0.3 cm/s) for CnF Vglut2-cre, 1.1 ± 0.04 s (at 12.2 ± 0.1 cm/s) for CnF VGAT-cre, 1.1 ± 0.03 s (at 12.5 ± 0.4 cm/s) for PPN Vglut2-cre, 1.1 ± 0.02 s (at 12.1 ± 0.3 cm/s) for PPN VGAT-cre, and 1.1 ± 0.05 s (at 13.4 ± 0.6 cm/s) for PPN ChAT-cre mice. In the 5 mouse groups, locomotor bout durations were similar (*P* > 0.05 one-way Kruskal-Wallis ANOVA on ranks) and locomotor speeds during these bouts were similar (11.9-13.4 cm/s, *P* > 0.05 one-way Kruskal-Wallis ANOVA on ranks) (see also Figure 4G-4L, bottom).

**Figure 4.**
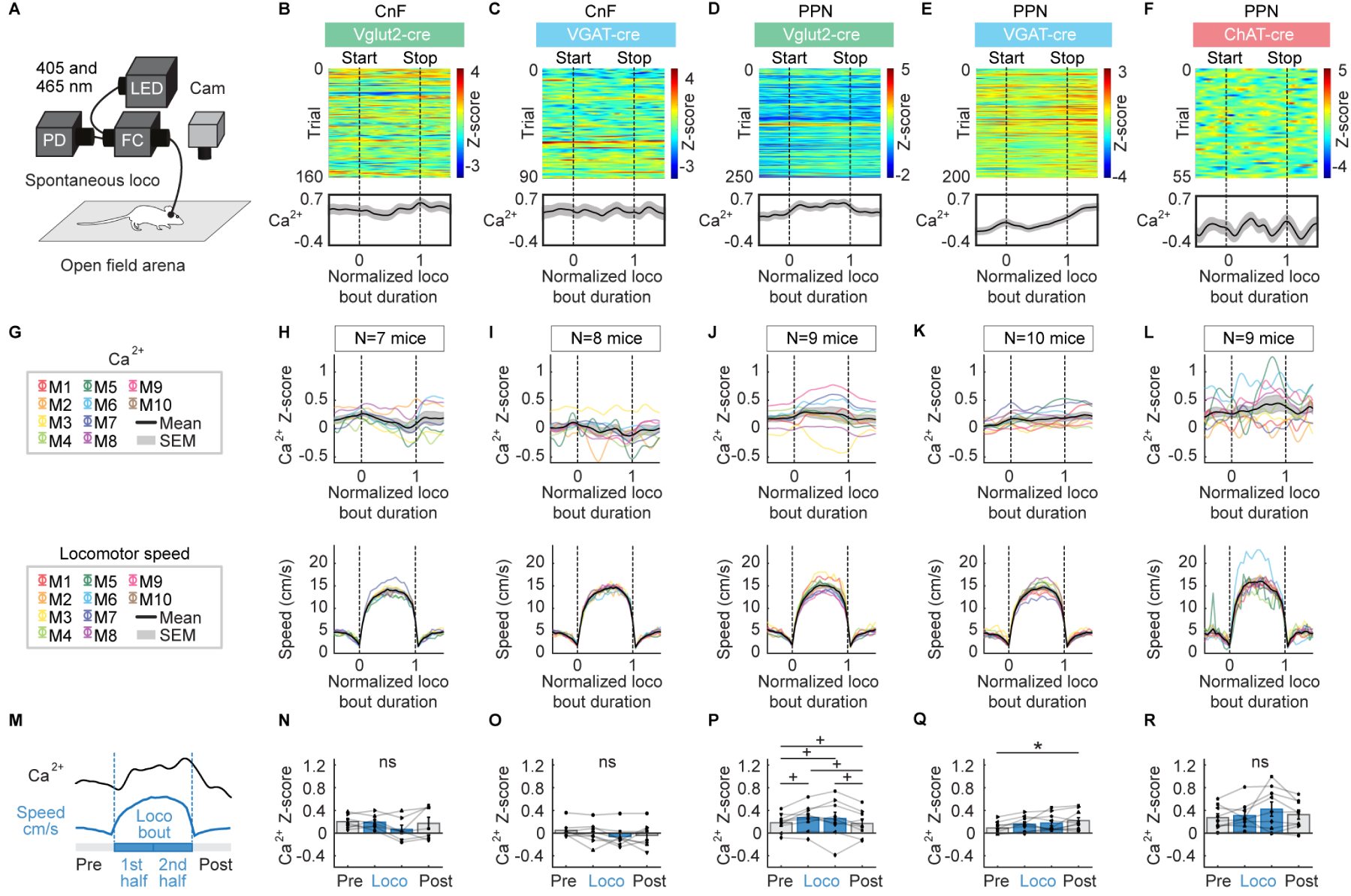
CnF or PPN cell activity during spontaneous, self-paced locomotion. (A) Fiber photometry recordings were done as in Figure 1. In an open-field arena, spontaneous locomotor bouts (speed > 3 cm/s for at least 0.5 s)^8,9^ were detected and aligned with calcium (**Ca^2+^**) signals recorded in the CnF or PPN. (B-F) Ca^2+^ activity in the CnF or PPN cells color-coded as in Figure 1 during spontaneous locomotion for example animals. Each line illustrates a locomotor bout. Locomotor bout onsets (0) and offsets (1) are indicated by black vertical dotted lines. Time is normalized as a fraction of locomotor bout duration. (G-L) Mean Ca^2+^ activity ± SEM (top) and locomotor speed ± SEM (bottom) recorded during spontaneous, self-paced locomotion in each mouse (M). The number of mice (N) used is indicated. (M-R) Mean Ca^2+^ signal before locomotion (-0.5 to 0), during the first half of the bout (0 to 0.5), second half of the bout (0.5 to 1) and after (1 to 1.5) locomotion (95-294 bouts for CnF Vglut2-cre, 34-343 bouts for CnF VGAT-cre mice, 113-328 bouts for PPN Vglut2-cre, 79-301 bouts for PPN VGAT-cre, 10-102 bouts for PPN ChAT-cre mice) (ns, not significant, \**P* < 0.05 Student-Newman-Keuls test after significant one-way repeated measures ANOVA; ^+^*P* < 0.05 Student Newman-Keuls test after a significant one-way repeated measures ANOVA on ranks).

We analysed the mean Ca^2+^ signal during four time periods: before the locomotor bout, during the first half, and second half of the locomotor bout, and after the locomotor bout (Figure 4M). In PPN Vglut2^+^ cells, the Ca^2+^ signal was significantly higher during the first and second halves of the locomotor bout compared to before the bout, and significantly lower after the bout compared to the second half of the bout (Figures 4D,4G,4J,4M,4P, n = 9 mice). In PPN VGAT^+^ cells, the Ca^2+^ signal was significantly higher after the bout than before the bout (Figures 4E,4K,4Q, n = 10 mice). In contrast, Ca^2+^ signal did not significantly change during locomotion compared with before locomotion in CnF Vglut2^+^ cells (Figures 4B,4H,4N, n = 7 mice), CnF VGAT^+^ cells (Figures 4C,4I,4O, n = 8 mice) or in PPN ChAT^+^ cells (Figures 4F,4L,4R, n = 9 mice).

### Ca^2+^ activity in CnF and PPN neurons during grooming or rearing

We recorded Ca^2+^ signals during episodes of grooming or rearing with the video captured from the side of the animal (Figure 5 and Figure 6). Grooming and rearing were manually scored and synchronized with the Ca^2+^ signals. Grooming bouts lasted 6.8 ± 0.9 s for CnF Vglut2-cre (n = 7 mice), 5.2 ± 0.4 s for CnF VGAT-cre (n = 7 mice), 7.2 ± 0.7 s for PPN Vglut2-cre (n = 9 mice), 5.7 ± 0.7 s for PPN VGAT- cre (n = 10 mice), and 8.5 ± 0.9 s for PPN ChAT-cre mice (n = 9 mice). Grooming bout durations were similar across most mouse groups, except for the PPN ChAT-cre mice, which exhibited longer grooming bouts than CnF VGAT-cre mice (*P* < 0.05, Student-Newman-Keuls test after a *P* < 0.05 one-way ANOVA). Rearing bouts lasted 4.0 ± 0.7 s for CnF Vglut2-cre (n = 7 mice), 3.8 ± 0.3 s for CnF VGAT-cre (n = 7 mice), 3.9 ± 0.2 s for PPN Vglut2-cre (n = 9 mice), 4.2 ± 0.4 s for PPN VGAT-cre (n = 10 mice), and 3.2 ± 0.3 s for PPN ChAT-cre mice (n = 9 mice). The rearing bout durations were similar across the five mouse groups (*P* > 0.05, Kruskal-Wallis ANOVA on ranks).

**Figure 5.**
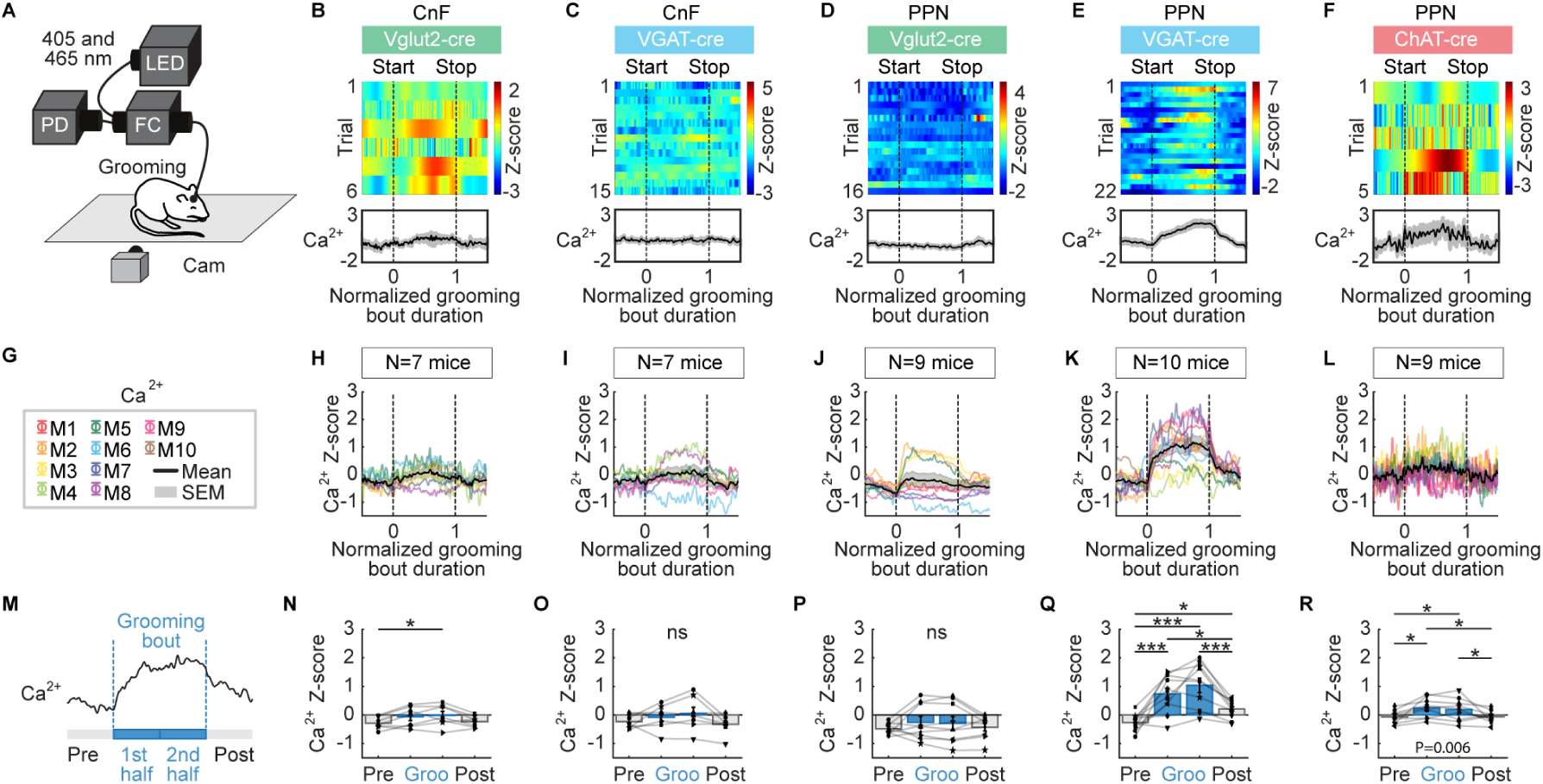
CnF or PPN cell activity during grooming. (A) Fiber photometry recordings were done as in Figure 1. Mice were filmed from the side on a treadmill that was switched off. Grooming bouts were detected and aligned with calcium (**Ca^2+^**) signals recorded in the CnF or PPN. (B-F) Ca^2+^ activity in the CnF or PPN cells color-coded as in Figure 1 during grooming for example animals. Each line illustrates a grooming event. Event onsets (0) and offsets (1) are indicated by black vertical dotted lines. Time is normalized as a fraction of the grooming event duration. (G-L) Mean Ca^2+^ activity ± SEM (top) recorded during grooming in each mouse (M). The number of mice (N) used is indicated. (M-R) Mean Ca^2+^ signal before grooming (-0.5 to 0), during the first half of the grooming bout (0 to 0.5), second half of the bout (0.5 to 1) and after grooming (1 to 1.5) (6-10 events for CnF Vglut2-cre, 3-15 events for CnF VGAT-cre mice, 7-39 events for PPN Vglut2-cre, 3-22 events for PPN VGAT-cre, 5-12 events for PPN ChAT-cre mice) (ns, not significant, \**P* < 0.05, \*\*\**P* <0.001, Student-Newman-Keuls test after a significant one-way repeated measures ANOVA).

**Figure 6.**
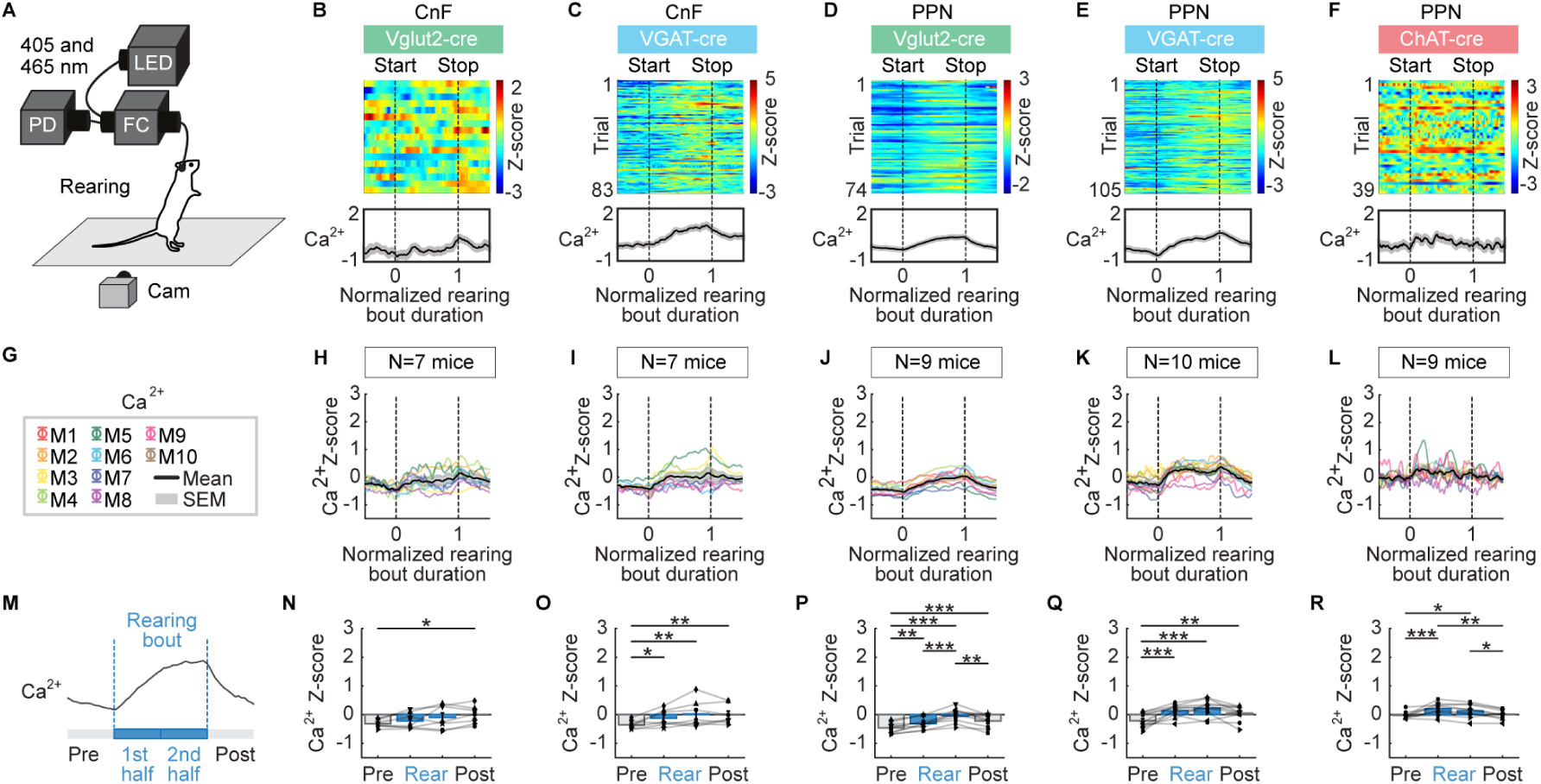
CnF or PPN cell activity during rearing. (A) Fiber photometry recordings were done as in Figure 1. Mice were filmed from the side on a treadmill that was switched off. Rearing bouts were detected and aligned with calcium (**Ca^2+^**) signals recorded in the CnF or PPN. (B-F) Ca^2+^ activity in the CnF or PPN cells color-coded as in Figure 1 during rearing for example animals. Each line illustrates a rearing event. Event onsets (0) and offsets (1) are indicated by black vertical dotted lines. Time is normalized as a fraction of the rearing event duration. (G-L) Mean Ca^2+^ activity ± SEM (top) recorded during rearing in each mouse (M). The number of mice (N) used is indicated. (M-R) Mean Ca^2+^ signal before rearing (-0.5 to 0), during the first half of the rearing bout (0 to 0.5), second half of the bout (0.5 to 1) and after rearing (1 to 1.5) (7-94 events for CnF Vglut2-cre, 17-86 events for CnF VGAT-cre, 13-79 events for PPN Vglut2-cre, 13-105 events for PPN VGAT-cre, 7-52 events for PPN ChAT-cre mice) (ns, not significant, \**P* < 0.05, \*\**P* < 0.01, \*\*\**P* < 0.001, Student-Newman-Keuls test after a significant one-way repeated measures ANOVA).

We analysed the mean Ca^2+^ signal during four time periods: before the grooming or rearing bout, during the first and second halves of the grooming or rearing bout, and after the grooming or rearing bout. During grooming behavior, PPN VGAT^+^ and PPN ChAT^+^ cells showed increased activity during the first and second halves of the bout compared to the pre-bout period, and a decreased Ca^2+^ signal after the bout compared to the second half (Figures 5A, 5E-5G, 5K-5M, 5Q-5R, n = 10 mice for PPN VGAT^+^ and n = 9 mice for PPN ChAT^+^ cells). CnF Vglut2^+^ cells displayed higher Ca^2+^ signal only during the second half of the grooming bout compared to the pre-bout period (Figures 5B, 5H, 5N). On the other hand, no significant change in Ca^2+^ signal was observed in CnF VGAT^+^ cells or PPN Vglut2^+^ cells during grooming compared to before the bout (Figures 5C-5D,5I-5J,5O-5P,5R, n = 7 mice for CnF VGAT^+^ and n = 9 mice for PPN Vglut2^+^ cells).

During the rearing behavior, VGAT^+^ cells in the CnF along with Vglut2^+^, VGAT^+^, and ChAT^+^ cells in the PPN showed an increase in Ca^2+^ signal during both the first and second halves of the bout compared to the pre-bout period, and PPN Vglut2^+^ and ChAT^+^ cells had a decrease in Ca^2+^ signal after the bout, relative to the second half of the rearing bout (Figures 6A, 6C-6F, 6G, 6I-6L, 6M, 6O-6Q, n = 7 to 10 mice per cell type). In contrast, CnF Vglut2^+^ cells only exhibited a higher Ca^2+^ signal after the bout compared to the pre-bout period (Figures 6B,6H,6N, n = 7 mice).

## DISCUSSION

In the present study, using fiber photometry, we recorded Vglut2^+^, VGAT^+^ or ChAT^+^ cells in the CnF or PPN during locomotion, non-locomotor behaviors, and following sensory stimulation (Figure 7). Vglut2^+^ and VGAT^+^ cells responded to sound, visual looming, or air puffs, except for PPN VGAT^+^ cells that did not respond to visual looming. PPN ChAT^+^ cells responded to air puffs. Air puffs triggered faster locomotor responses (∼30-35 cm/s) than sound or visual looming (∼10-20 cm/s). When high-speed locomotion was triggered by an air puff, activity increased in CnF Vglut2^+^ neurons. When low-speed locomotion was triggered by sound or visual looming, activity increased in PPN Vglut2^+^ neurons. During self-paced locomotion (low speed, ∼10-20 cm/s), activity increased in PPN Vglut2^+^ cells, while in PPN VGAT^+^ cells, activity was higher after the bout compared with before. During grooming, activity increased in CnF Vglut2^+^ cells, PPN VGAT^+^ and ChAT^+^ cells. During rearing, activity increased in CnF VGAT^+^ cells and PPN Vglut2^+^, VGAT^+^, and ChAT^+^ cells. Altogether, our study reveals cell type-specific activity in the CnF or PPN during locomotor and non-locomotor behaviors, or following sensory stimulations. Such responsiveness prompts questions about how sensory information shapes locomotor commands in the CnF and PPN. Our results may have clinical relevance, as sensory circuits are increasingly being targeted to improve locomotor function in pathological states such as Parkinson’s disease.

**Figure 7.**
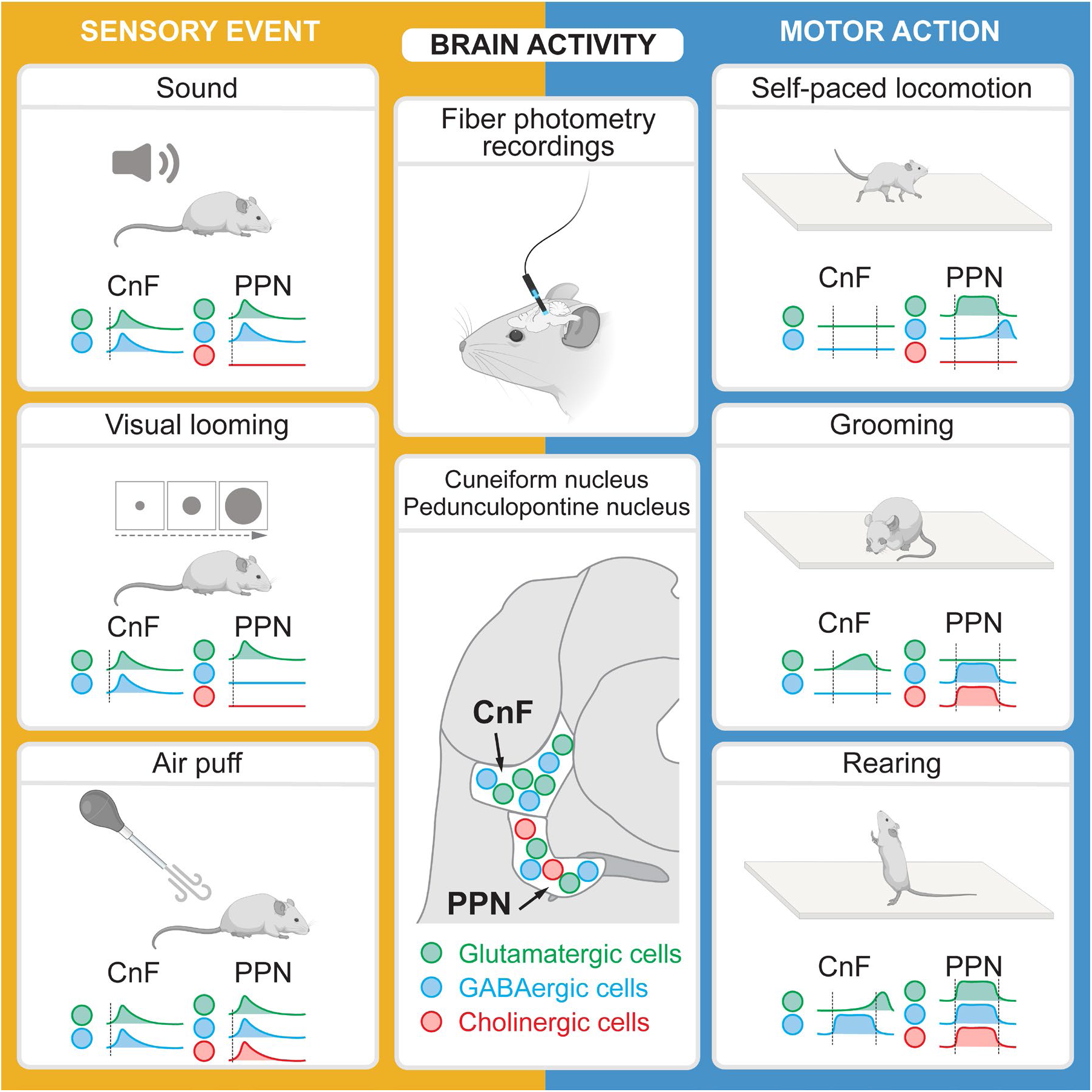
Sensory and motor-related activities in genetically-defined cell types in the CnF and PPN. On the left, for each stimulus, the statistically significant increases in activity per cell population are illustrated. The vertical dotted lines indicate the timing of sensory stimulation. On the right, for each motor behavior, the statistically significant increases in activity per cell population are shown. The vertical dotted lines represent the beginning and the end of the motor bout.

### CnF or PPN activity during motor behaviors

During self-paced locomotion, we found that activity increased in PPN Vglut2^+^ neurons. This is in accordance with the key role of MLR glutamatergic neurons in the generation of the locomotor drive sent to reticulospinal neurons from lamprey to mammal (e.g. lamprey^29^, mouse^31^, for review^3^). The selective recruitment of PPN Vglut2^+^ neurons (but not CnF Vglut2^+^ neurons) at the slow speeds exhibited by our mice in the open field (∼10-20 cm/s) aligns with findings showing that optogenetic activation of PPN Vglut2^+^ neurons selectively evokes slow locomotion during exploration^5,11^. Our findings are also consistent with electrophysiological recordings showing that the firing of glutamatergic PPN neurons positively correlates with locomotor speed (PPN Vglut2^+^ neurons^5^, Vglut2^+^ neurons without distinction between CnF and PPN^13^, putatively glutamatergic CamKIIa^+^ without distinction between CnF and PPN^16^, CamKIIa^+^ PPN neurons^17^, unidentified PPN neurons^18^). During air puff-evoked, high-speed locomotion (∼30-35 cm/s), we found increased activity in CnF Vglut2^+^ neurons, consistent with their role in producing the fastest gaits (gallop and bound) during air puff-evoked escape behavior, as demonstrated by chemogenetic inhibition^5^.

The PPN not only contains Vglut2^+^ cells that induce locomotion. It is a functionally heterogeneous area which also includes some Vglut2^+^ cells that stop locomotion (for review^4^). Our recordings of bulk fluorescence with fiber photometry thus reflects activity from intermingled glutamatergic populations. We probably recorded a combination of predominantly pro-locomotor Vglut2^+^ PPN neurons (i.e. whose optogenetic activation initiates locomotion^5,11^, see also^13,31,16,17^ and a much lesser contribution of glutamatergic PPN neurons that stop locomotion^6,7^, see also Chx10^+^ PPN neurons^32^; Rbp-4-transgene^+^ neurons^20^; CamKIIa^+^ neurons^17,33^). If few PPN glutamatergic stop cells were recorded, their activity may have been masked by averaging. Single-cell recordings would be needed to reveal stop-related activity among Vglut2^+^ cells. Future studies should also examine whether stop-related signals are linked to Chx10^+^ cells in rostral PPN^32^, and locomotion signals to Vglut2^+^ neurons in caudal PPN^5,11^. Finally, part of the activity recorded during locomotion in the PPN could also result from sensory feedback generated during locomotion. Indeed, spiking changes in unidentified PPN cells during limb movements in cats^34^, during passive and active limb movements in monkeys^35,36^ and in humans^37–39^.

During rearing we observed increased activity in all CnF and PPN cell types except the Vglut2^+^ cells of the CnF, which showed increased activity only after the rearing bout. The PPN Vglut2^+^ cells that were active during rearing likely included spinal cord-projecting neurons known to encode rearing and to control body extension^20^. During grooming, we observed increased activity in CnF Vglut2^+^ cells, PPN VGAT^+^ and PPN ChAT^+^ cells. We found no significant increase in activity of PPN Vglut2^+^ cells, even though putatively glutamatergic PPN neurons expressing the Rbp-4-transgene encode handling and grooming^20^. A possible explanation is that the PPN also contains glutamatergic neurons expressing Chx10, which stop grooming and rearing behavior when stimulated optogenetically^32^. On top of that, the Rbp-4^+^ neurons that encode grooming and handling are predominantly located in the mesencephalic reticular formation, which medially borders the CnF and PPN, with a relatively lower number of Rbp-4^+^ neurons located in the PPN^20^. The VGAT^+^ neurons we recorded during grooming or rearing may be local interneurons that prevent pro-locomotor cells from being recruited^13,5^. In line with this possibility, we recorded increased activity in PPN VGAT^+^ cells after the locomotor bouts. Future studies should use single-cell recordings to determine if the same CnF or PPN cells are active during locomotor bouts, locomotor stops, rearing, and grooming.

### Sensory-evoked activity in CnF and PPN cells

Regarding sound stimulations, our results align with previous studies showing sound-evoked electrophysiological responses in unidentified CnF and PPN neurons in cats^40,41,34^, rats^42^ and monkeys^43^, and calcium responses in PPN CamKIIa^+^ neurons in mice^44^. A possible source of inputs to the cells we recorded could be the inferior colliculus which processes auditory information (for review^45,46^) and projects to CnF and PPN glutamatergic neurons in mice^13,5,7^.

We did not measure whether sound stimulation evoked an acoustic startle reflex, i.e. a rapid recoil of the head and ears, hunching of the back, and extension of the limbs and tail^47,48^. To accurately capture this rapid reflex which occurs 10-200 ms after sound onset, high-speed recordings (1,000 frames/s) and/or a piezoelectric startle platform are required^49,48^. Our sound stimulation evoked a low proportion of locomotor responses (12-44% trials). This may be related to the properties of the sound used here, particularly its short duration (∼0.5 s Figure S2A,2D). Longer duration sounds (5 s, 70-80 dB white noise) appear to evoke escape more frequently^50,51^.

Regarding visual looming, previous studies have reported light flash-evoked electrophysiological responses in unidentified CnF and PPN neurons in rats^42^ and monkeys^43^, suggesting that visual information is encoded in such cells. However, such neutral stimuli do not carry any valence in contrast with a visual looming stimulus, which likely also recruits circuits beyond visual processing. Visual looming is known in mice to trigger flight or freezing responses through the involvement of threat-processing regions such as the superior colliculus (mice^52^, see also in lamprey^53^) or periaqueductal gray^52,54^ that both project to the CnF and _PPN13,5,7._

The disk expansion rates we used encompassed the range previously tested for visual looming stimuli that trigger escape responses in mice (35-350 degrees/s^55^). In our data, the proportion of locomotor responses evoked by visual looming was low (13-27% trials), and this could be due to several factors. First, mice exhibit reduced escape behavior when no shelter is present in the arena^56^. Second, the effect of visual looming likely depends on the mouse’s position relative to the screen as they have a 280° field of vision^57^. Third, mice may have learned to predict the visual looming, as due to our synchronization procedure (see Methods), visual looming was preceded by a button press that generated a small noise (58.6 ± 0.5 dB, Figure S2C,S2F), which in some mice elicited a slight increase in the photometry signal (Figures 2H-2L). Finally, it is worth to note that we did not quantify whether the non-locomotor responses corresponded to freezing.

Regarding air puffs, our results are in line with air puff-evoked electrophysiological responses recorded in Vglut2^+^ neurons without distinction between CnF and PPN^58^. Importantly, an air puff is a multifaceted stimulus involving a somatosensory component, the sound of the puff, and a visual looming caused by the hand bringing the air bulb close to the mouse’s back. The somatosensory component likely involves the ascending sensory pathways^59^. However, it is likely that again, threat-processing regions were recruited such as amygdala (mouse^58^, monkey^60^) which projects to the CnF/PPN region, and whose manipulation influences the amplitude of air-puff evoked responses in CnF/PPN cells^58^.

The habituation that we observed in some CnF or PPN neurons could indicate that these neurons encode the saliency of sensory events. Single-cell recordings suggest that some cells in the PPN or CnF respond to multiple sensory modalities and therefore rather encode stimulus saliency^42,43^. The PPN is part of the reticular activating system, which regulates arousal towards salient sensory events, i.e. carrying behaviorally relevant information^61–67^ (for review^68,10^). PPN neurons are considered to serve as an interface between sensory and motivational systems, and were reported to provide reward-, sensorimotor- and arousal-related signals to other brain regions^34,69,70,42,43^ (for review^10^). As reported here, some PPN and CnF cells are activated by sensory stimuli and display habituation to sound stimuli as in monkey^43^. Other authors have proposed that PPN cells encode the saliency of current events in comparison to anticipated ones, and thereby contribute to regulate behavioral flexibility^71^ (see also^10^).

### A balanced network for competition between behavioral outputs?

A striking observation in the present study is that Vglut2^+^ and VGAT^+^ cells in the CnF or PPN were recruited simultaneously by most sensory stimuli. This is reminiscent of the almost simultaneous increase of excitatory and inhibitory synaptic conductances evoked by a sound in auditory cortical neurons (rat^72,73^; mouse^74^) or by deflection of a whisker in somatosensory cortical neurons^75^. Our data suggest the existence in the CnF and/or PPN of a balanced network where glutamatergic and GABAergic neurons compete for behavioral output. Populations of glutamatergic cells driving distinct outputs would be recruited by incoming inputs, and competitive inhibition could play a gating role, as in other brain regions^76–79^ (for review^80–82^). In support of this possibility, PPN Vglut2^+^ cell activity was higher during sound- or visual looming-evoked locomotion, and CnF Vglut2^+^ cell activity was higher during air puff-evoked locomotion. In trials where sensory stimulation did not elicit locomotion, sensory-evoked CnF Vglut2^+^ cell activity was probably not strong enough to counteract concurrent activation of CnF VGAT^+^ neurons, which send GABAergic input to local VGAT-negative neurons (i.e. presumably CnF Vglut2^+^ ones^13,5^). Alternatively, the absence of locomotor response could involve the recruitment of other cells inhibiting locomotion such as PPN Chx10^+^ neurons^32^, or cells downstream of the MLR such as reticular Chx10^+^ neurons in the gigantocellularis nucleus^83,84^. A balanced network organisation in the PPN could also be at play in the context of signaled active avoidance, where a sound predicting threat is conditioned to elicit locomotion directed to a safe zone^85^. In line with such organization, blockade of PPN putative glutamatergic (CamKIIa^+^) neurons decreases active avoidance, whereas blockade of PPN VGAT^+^ neurons facilitates active avoidance^85^.

### Clinical relevance

In Parkinson’s disease, sensory circuits are increasingly targeted to improve locomotor function. To compensate for the deficit of internal cueing and/or proprioceptive integration, clinicians use cueing with visual stimuli (stripes on the floor^86^, for review^87^) sound stimuli (rhythmic sounds^86^, for review^88^; music^89^) or somatosensory stimuli (vibration applied to wrist^90^; or to lower leg^91^) (for meta-analyses^92,24,25^). However, sensory inputs can also worsen locomotor performance, as visual information such as passing a door can evoke gait freezing^87^. It is still unclear which circuits are involved during cue-assisted locomotion and sensory-evoked freezing. Our recordings indicate that during sensory-evoked locomotion, activity was higher in Vglut2^+^ neurons of the PPN or CnF. Tailoring interventions to specifically activate some of these cells may favor locomotor activity. In line with this, selective activation of Vglut2^+^ neurons in the CnF or Vglut2^+^ neurons in the caudal PPN promotes locomotion in rodent models of Parkinson’s disease, without interfering with the ability to adapt navigation^9,11^.

### Limitations of the study

Fiber photometry does not allow us to determine whether the same cells that control locomotion (so-called “MLR” cells) also encode sensory information. It is possible that distinct intermingled cells are activated during locomotion or sensory stimulation. Nevertheless, other elements of the locomotor circuitry are known to encode both locomotor commands and sensory information, such as reticulospinal neurons (e.g. lamprey^93–95^; salamander^96^; mouse^97^, for review^98^) or spinal cells of the central pattern generator (lamprey: for review^99^; zebrafish^100,101^). Future studies should address this issue using single-cell recordings.

Regarding photometry, we cannot exclude the possibility of partial contamination from PPN cells during recordings in the CnF, and vice versa, due to their close proximity, as well as potential contamination by adjacent areas such as the periaqueductal gray, inferior colliculus, or mesencephalic reticular formation. Based on modeling and measurements made in brain tissue, we can estimate that the recorded Ca^2+^ signals should originate from a maximum of ∼500 µm below fiber tip^102,103^.

## Methods

### Ethics statement

All procedures were in accordance with the guidelines of the Canadian Council on Animal Care and were approved by the animal care and use committees of the Université de Sherbrooke (QC, Canada). Care was taken to minimize the number of animals used and their suffering.

### Animals

Homozygous Vglut2-cre knock-in mice^27^, VGAT-cre knock-in mice^27^, and ChAT-cre knock-in mice^28^ were used. Vglut2-cre and VGAT-cre mice were bred in the laboratory and genotyped as previously described^104,8,9^. Homozygous ChAT-cre mice were obtained directly from Jax a few weeks before the experiments. Animals had ad libitum access to food and water, with lights on from 6 AM to 8 PM. Vglut2-cre mice used were 16-24 weeks old at time of use (10 males, 7 females). VGAT-cre mice used were 15-30 weeks old at time of use (10 males, 9 females). ChAT-cre mice used were 10-12 weeks old at time of use (5 males, 4 females).

### Virus injection and optic fiber implantation

The procedure was adapted from previously reported ones^105,8,9,29,106^. Briefly, mice were anaesthetized using isoflurane (induction: 5%, 500 mL/min; maintenance: 1.5–2.5%, 100 mL/min) delivered with a SomnoSuite (Kent Scientific, Torrington, CT, USA). Mice were placed in a Robot Stereotaxic instrument coupled with StereoDrive software (Neurostar, Tübingen, Germany). Mice were given buprenorphine as an analgesic (0.1mg/kg s.c., volume 0.3 mL). An incision was made on the scalp, a hole was drilled in the cranium and a 10 µL Hamilton syringe locked into the Robot Stereotaxic instrument was used to inject unilaterally in CnF or in PPN a volume of 300 nL^5^ of a solution containing an adeno-associated virus (AAV) driving the expression of a calcium sensor jGCaMP7f in a cre-dependent manner (pGP-AAV-syn-FLEX-jGCaMP7f-WPRE (Addgene plasmid # 104492; RRID:Addgene_104492), titer 7×10^12^ particles per milliliter). The AAV solution was injected unilaterally into the CnF (anteroposterior -4.80 mm, mediolateral +1.10 to mm, dorsoventral -2.90 mm relative to bregma) or the PPN (anteroposterior -4.72 mm, mediolateral +1.20 mm, dorsoventral -3.75 mm relative to bregma) at a rate of 0.05 µL/min. The syringe was left in place for 1 min before being removed. Then, an optic fiber (400 µm core, 0.66 NA, 3 to 5 mm length, Doric Lenses) held in a stainless-steel ferrule was placed 150 µm above the right CnF at -4.80 mm anteroposterior, +1.10 mm mediolateral, -2.75 mm dorsoventral or right PPN at - 4.72 mm mm anteroposterior, +1.20 mm mediolateral, -3.60 mm dorsoventral relative to bregma^8,9^. The ferrule was secured on the cranium using two 00-96×1/16 mounting screws (HRS Scientific, QC, Canada) and dental cement (A-M Systems, Sequim, WA, USA).

### Fiber photometry recordings and analyses

For excitation of the jGCaMP7f, two LEDs (Doric Lenses) sent light with a wavelength of 465 nm for calcium-dependent fluorescence and 405 nm as an isosbestic control with light intensity sinusoidally modulated at respectively 211Hz and 531Hz^29^. The light from the two LEDs was sent through a filter cube (FMC4, excitation wavelength range 400-410 nm and 460-490 nm, emission wavelength range 500-550 nm, Doric Lenses) which converges it into a fiber optic patch cord (low auto-fluorescence MFP, 400 µm core, 0.57 NA, Doric lenses) with rotary joint (FRJ, 400 µm core, 0.57 NA Doric Lenses) attached to a fiber optic cannula implanted in the mouse brain (see section above). jGCaMP7f fluorescence was captured by the fiber optic cannula, sent back through the fiber optic patch cord, and directed to a photoreceiver (Visible Femtowatt Photoreceiver Model 2151, Newport) by the filter cube. Lastly the fluorescence detected by the photoreceiver was demodulated by a lock-in amplifier and data acquisition system (RZ5P; Tucker Davis Technologies). The fiber photometry system was controlled through a PC with Synapse Suite software (Tucker Davis Technologies).

For the analysis, a custom Matlab script^29^ was used to perform a Fourier transformed subtraction of the photometry signals and a subsequent smoothing with a high-pass Butterworth filter (3rd order, cutoff frequency 0.0051 Hz) and a low-pass Butterworth filter (5th order, cutoff frequency 2.29 Hz). Subtracted and smoothened signals were then normalized per recording as a Z-score (Z-score = (signal - mean) / SD). To analyze the changes in Ca^2+^ Z-scores evoked by sensory stimulations, raw time windows (from -5 s to 10 s after stimulation) were extracted for each trial and the recorded signals (photometry signal and locomotor speed) were averaged, with locomotor speed first binned in 50 ms time bins. To analyze the changes in Ca^2+^ Z-scores during multiple occurrences of a specific of motor event (such as a self-paced locomotor bout, a grooming or a rearing bout) which systematically vary in duration, we normalized time as a function of the duration of the motor event on a scale from 0 to 1. Then, normalized time windows (-0.5 to 1.5 of normalized time) were extracted for each motor event, and the corresponding photometry signal and motor parameter (such as locomotor speed) were averaged per normalized time bin, which was defined as 0.2 % of normalized time for the photometry signal and 5 % of normalized time for the motor parameter.

### Open-field locomotion

The procedure was as previously reported^8,9,106^. Briefly, locomotor activity was filmed from above in a 40 × 40 cm open-field arena at 20 fps using a Logitech Brio camera. Locomotor activity was recorded during trials of 10 min. Video recordings were analyzed with DeepLabCut to track user-defined body parts^107–109^ and a custom Matlab script (Mathworks, Natick, MA, USA)^8,9,106^. We tracked the body center positions, the corners of the arena for distance calibration, and the small LED to detect optogenetic stimulations. Timestamps were extracted using Video Frame Time Stamps (Matlab File Exchange). Body center positions and timestamps were used to calculate locomotor speed. Body center and tail base positions were excluded if their detection likelihood by DeepLabCut was <0.8, if they were outside of the open-field arena, or if body center speed exceeded the maximum locomotor speed recorded in mice (334 cm/s^110^). A locomotor bout was defined when body center speed was higher than 3 cm/s for at least 0.5 s^8,9^.

### Rearing and grooming

Mice were placed on a 37.5 × 5.1 cm treadmill (Model 8709, Letica Scientific Instruments, Panlab, Spain) and speed was set at 0 cm/s. Animal behavior was filmed from the side at 20 fps using a Logitech Brio camera. Onsets and offsets of rearing and grooming bouts were manually scored during visual examination of the video and synchronized with the fiber photometry recordings offline.

### Sensory stimulations

For all stimulations, mice were filmed from above in an open-field arena. *Sound stimulation.* A clinking metallic sound (86.9 ± 0.5 dB, n = 5 trials, sound spectrogram illustrated in Figures S2A,S2B) was applied every 60.1 s ± 0.3 s, ∼1 m away from the open-field arena. *Visual looming stimulation.* A black disk increasing in diameter during 1.676 s on a grey background (disk expansion rates provided in Figures S3A-S3C) was shown every 61.1 ± 0.5 s on a 37.5 × 30 cm screen located laterally to the open field. *Air puff stimulation.* Air puffs were applied with a small air bulb on the back of the animal every 61.4 s ± 0.7 s. Such air puffs generated a noise (55.2 ± 0.5 dB, n = 5 trials, sound spectrogram illustrated in Figures S2B,S2D). To synchronize sensory stimulations with photometry, we used a button press on a Grass S88X stimulator, located 20 cm away from open-field arena, that sent a TTL signal to the acquisition software. This button generated a small noise (58.6 ± 0.5 dB, n = 5 trials, sound spectrogram illustrated in Figures S2C,S2F). Sound intensities were measured ∼10 cm from the source with an iPhone 11 running Decibel X sound level meter software. Raw audio signal was imported into Matlab using the audioread function and sound spectrograms were generated using the spectrogram function.

### DeepLabCut networks

The networks used were the same as those described in^8,9,106^. Briefly, for the analysis of locomotion in the open-field arena, we labeled the body center, the tail base, the corners of the arena. We used a ResNet-50-based neural network^111,112^ with default parameters for 1,030,000 training iterations. We validated with one shuffle and found that the test error was 2.28 pixels and the train error 1.85 pixels^8,9,106^.

### Histology

Procedures were as previously reported^104,8,9,106^. Briefly, mice were anaesthetized using isoflurane (5%, 2.5 L per minute) and transcardially perfused with 30-50 mL of a phosphate buffer solution (0.1M) containing 0.9% of NaCl (PBS, pH = 7.4), followed by 50 mL of PBS solution containing 4% (wt/vol) of paraformaldehyde (PFA 4%). Post-fixation of the brains was performed for 24 h in a solution of PFA 4%. Brains were incubated in a PB solution containing 20% (wt/vol) sucrose for 24 h before histology. Brains were snap frozen in methylbutane (-45°C ± 5°C) and sectioned at -20°C in 40 µm-thick coronal slices using a cryostat (Leica CM 1860 UV). Floating sections at the level of the CnF and PPN were collected under a Stemi 305 stereomicroscope (Zeiss) and identified using the mouse brain atlas of Franklin and Paxinos^113^.

### Immunofluorescence

The procedure was as previously reported^104,8,9,106^. All steps were carried out at room temperature unless stated otherwise. The sections were rinsed three times during 10 min in PBS and incubated during 1h in a blocking solution containing 5% (vol/vol) of normal donkey serum and 0.3% Triton X-100 in PBS. The sections were incubated during 48 h at 4°C in a blocking solution containing a primary antibody against ChAT (ChAT; goat anti-choline acetyltransferase, Sigma AB144P, lot 3018862, 3675895, 4146422 (1:100), RRID: AB_2079751) and/or containing a primary antibody against green fluorescent protein (GFP) to label jGCaMP7f (chicken anti-GFP, Abcam, AB13970, lot GR3190550-23, GR3190550-30, GR3361051-12, GR1018753-22 (dilution 1:500), RRID: AB_300798) and gently agitated with an orbital shaker. Then, the sections were washed three times in PBS and incubated during 4 h in a blocking solution containing a secondary antibody to reveal ChAT (donkey anti-goat Alexa 594, Invitrogen A11058, lot 1975275, 2306782 (1:400), RRID: AB_2534105) or GFP (donkey anti-chicken Alexa 488, Jackson ImmunoResearch 703-545-155, lot 135111,154923 (1:400), RRID: AB_2340375). The slices were rinsed three times in PBS for 10 min and mounted on Colorfrost Plus slides (Fisher 1255017) with a medium with DAPI (Vectashield H-1200) covered with a 1.5 type glass coverslip and stored at 4°C before observation.

### Microscopy

Brain sections were observed using a Zeiss AxioImager M2 microscope bundled with StereoInvestigator 2018 software (v1.1, MBF Bioscience). Composite images were assembled, and the levels were uniformly adjusted in Photoshop CS6 (Adobe) to make all fluorophores visible and avoid pixel saturation, and digital images were merged.

### Statistics and reproducibility

Data are presented as mean ± standard error of the mean (SEM) unless stated otherwise. No statistical method was used to pre-determine sample sizes, which are similar to those used in the field (e.g.^5,8^). Statistical details of experiments can be found in the figure legends, including the sample size and the number of trials recorded per animal. No randomization or blinding procedure was used. Statistical analysis was done using Sigma Plot 12.0. Parametric analyses were used when assumptions for normality and equal variance were respected, otherwise non-parametric analyses were used. Normality was assessed using the Shapiro-Wilk test. Equal variance was assessed using the Levene test. To compare the means between two dependent groups, a parametric two-tailed paired t-test or a non-parametric Wilcoxon test was used. To compare the means between two independent groups, a two-tailed t-test or a non-parametric Mann-Whitney rank sum test was used. For more than two dependent groups, a parametric one-way analysis of variance (ANOVA) for repeated measures or a non-parametric Friedman ANOVA on ranks was used. For more than two independent groups, a parametric one-way ANOVA or a non-parametric Kruskal-Wallis ANOVA on ranks was used. ANOVAs were followed by a Student Newman-Keuls post hoc test for multiple comparisons between groups. Linear or nonlinear regressions between variables and their significance were calculated using Sigma Plot 12.0. Statistical differences were assumed to be significant when *P* < 0.05.

## DATA AVAILABILITY

The datasets generated during and/or analysed during the current study are available from the corresponding author on reasonable request.

## CODE AVAILABILITY

This paper does not report original code.

## ACKNOWLEDGMENTS

This work was supported by the Canadian Institutes of Health Research (407083 and 506880 to D.R.); the Natural Sciences and Engineering Research Council of Canada (RGPIN-2017-05522, RTI-2019-00628 and RGPIN-2024-05567 to D.R.); the Fonds de la Recherche du Québec - Santé (FRQS Junior 1 awards 34920 and 36772, and Junior 2 award 297238 to D.R.; Doctoral training scholarship 335559 to C.I.v.d.Z.); the Canada Foundation for Innovation (39344 to D.R.), the Centre de Recherche du Centre Hospitalier Universitaire de Sherbrooke (start-up funding and PAFI grant to D.R.), the Faculté de médecine et des sciences de la santé (start-up funding to D.R.), the Centre d’excellence en Neurosciences de l’Université de Sherbooke (to D.R.), and the fonds Jean-Luc Mongrain de la Fondation du CHUS (to D.R.). This study has received funding from the European Research Council (ERC) under the European Union’s Horizon 2020 research and innovation programme (grant agreement No 951477 to D.R.). We thank Pr. Gendron for providing access to the stimulator, Pr. Sarret for providing access to the stereotaxic robot and cryostat, Pr. Denault for providing access to a -80° freezer, Véronique Blais for technical assistance at an early stage of this study, and Pr. Pisanello for helping us evaluate the volume of collected light beneath the fiber. We thank Rachel Donka for the discussion on photometry data analysis.

## AUTHOR CONTRIBUTIONS

CIvdZ: conceptualization, data curation, formal analysis, investigation, methodology, software, validation, visualization, writing—original draft, writing— review and editing. AJT, JSS, JB, JDY: data curation, investigation, methodology, writing—review and editing. TH, VK, MFR: methodology, writing—review and editing. DR: conceptualization, data curation, formal analysis, funding acquisition, methodology, project administration, resources, supervision, validation, visualization, writing—original draft, writing—review and editing. All authors contributed to the article and approved the submitted version.

## SUPPLEMENTARY INFORMATION

**Figure S1.**
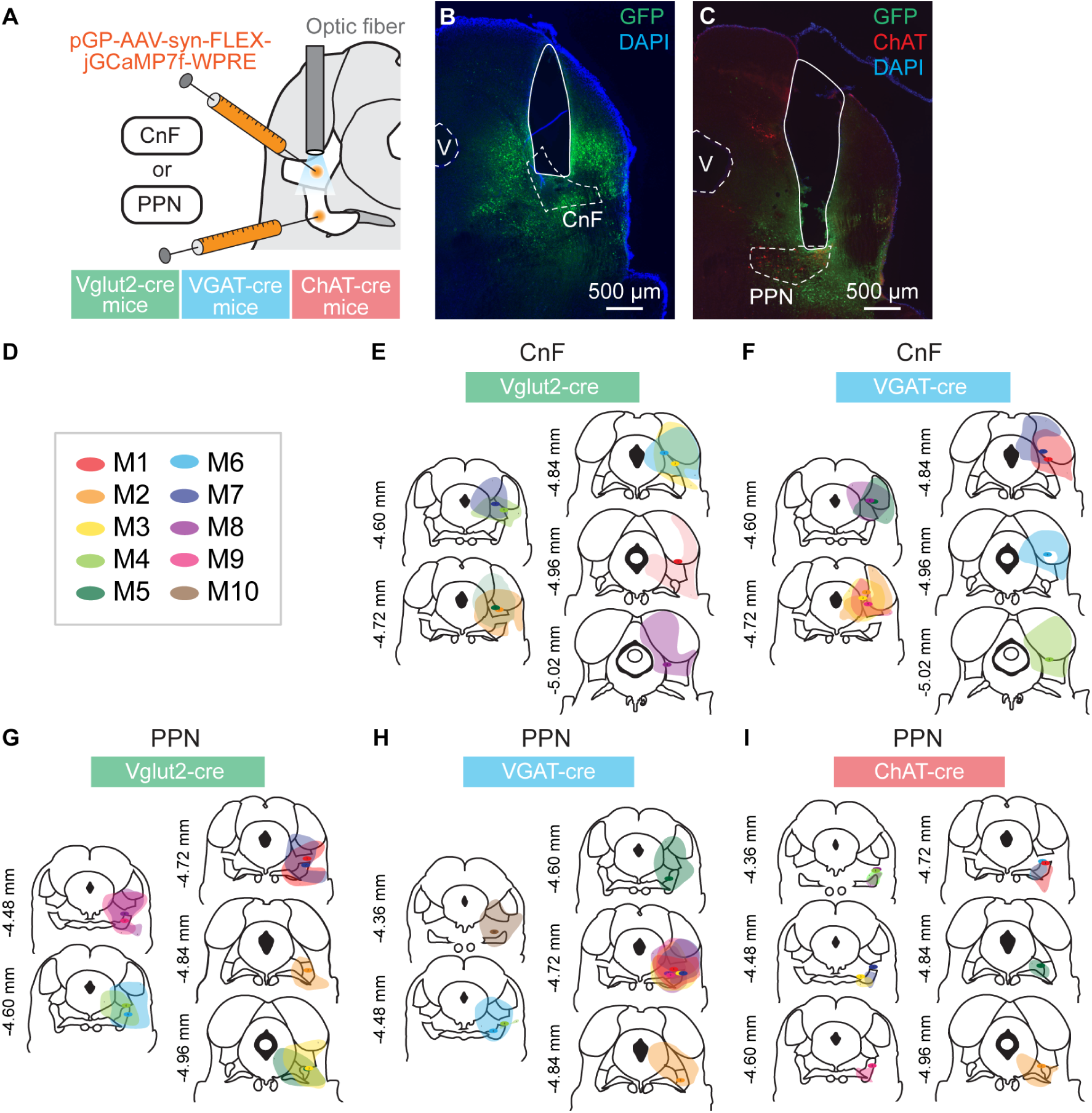
Adeno-associated virus injections and photometry fiber implantations. **(A)** Vglut2-cre, VGAT-cre or ChAT-cre mice were injected in the cuneiform nucleus (**CnF**) or pedunculopontine nucleus (**PPN**) with an adeno-associated virus (**AAV**) encoding for the genetically encoded calcium indicator jGCaMP7f in a cre-dependent manner (see Methods) and implanted with an optic fiber ∼150 μm above the injection site. **(B,C)** Photomicrographs showing the position of cells infected by virus injection (green) in CnF or PPN and the position of the optic fiber right above the CnF (B) or PPN (C) in example Vglut2-cre mice, with the nuclear marker DAPI shown in blue. In C, cells immunoreactive for choline acetyltransferase (**ChAT**) are shown in red. **(D)** Color code used to distinguish individual mice (M). **(E-I)** Histological locations of the cells infected by the AAV and of the tips of optic fibers.

**Figure S2.**
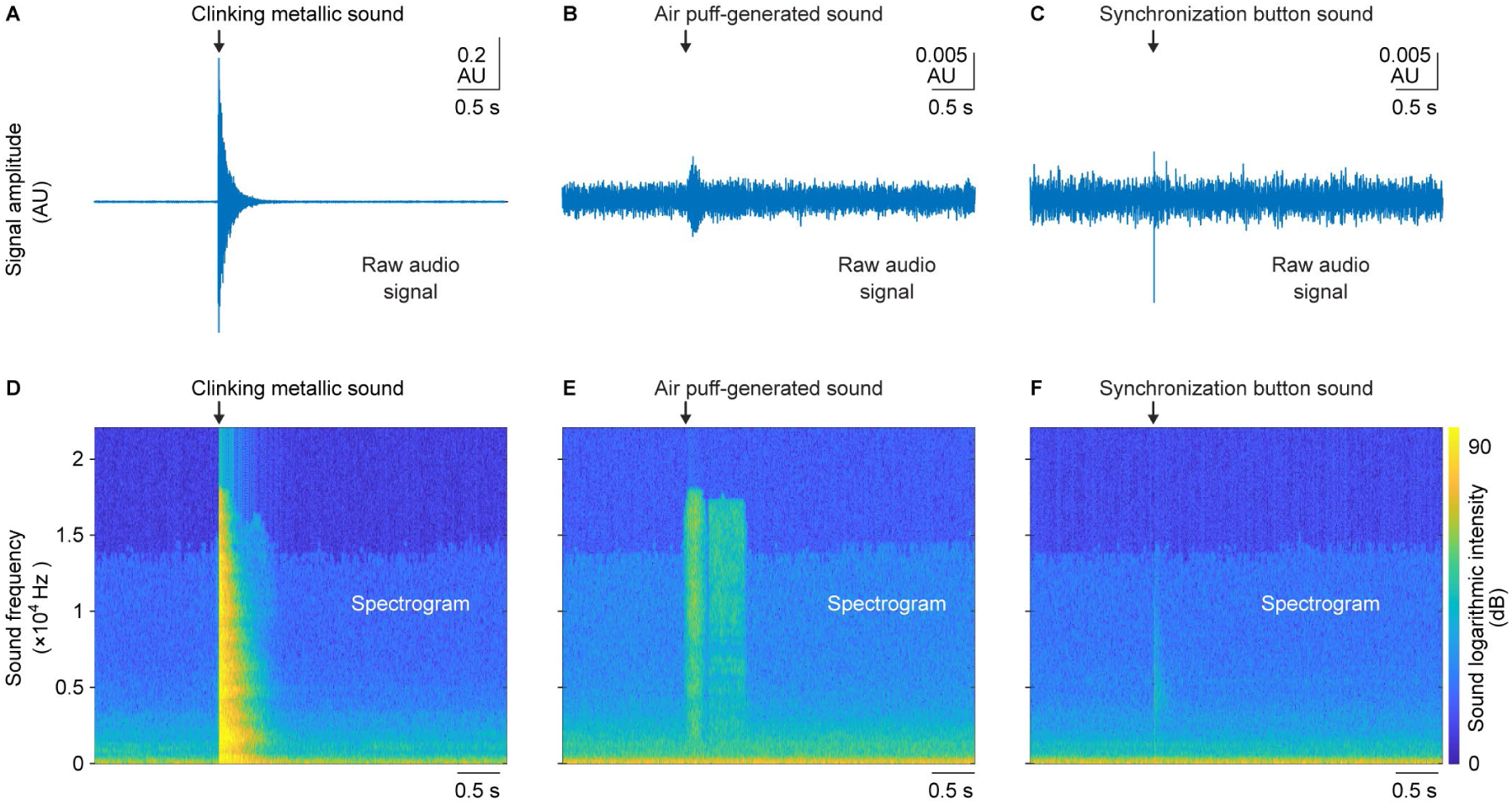
Sound spectrograms. **(A-C)** Raw audio signal for a clinking metallic sound (A), air puff-generated sound (B), and for the sound generated by the button press used to synchronize sensory stimulation and fiber photometry recordings (C). These signals were used to generated spectrograms in D-F (see Methods). **(D-F)** Typical spectrograms obtained for a clinking metallic sound (D), air puff-generated sound (E), and button press sound (F). The x-axis represents time, with a scale similar to that used for raw audio signals in (A-C). The y-axis represents sound frequencies, and the color scale indicates the logarithmic intensity of the sound.

**Figure S3.**
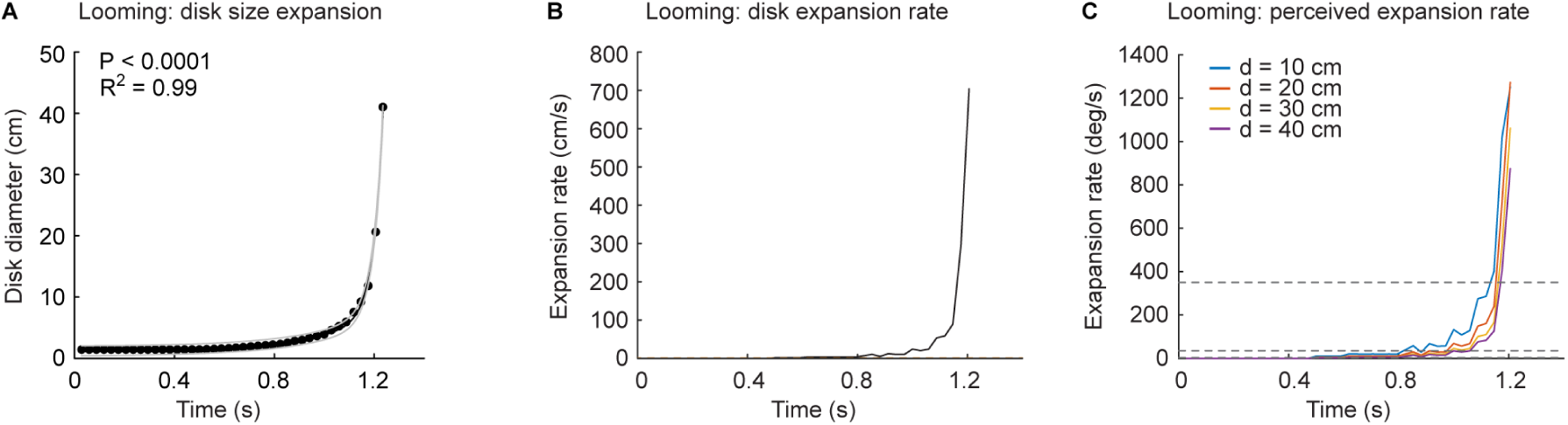
Visual looming stimulus properties. **(A)** Black disk diameter grew exponentially as a function of time throughout the duration of the visual looming stimulus (∼1.7 s). Note that beyond 1.2 s, part of the disk exceeds the screen size, making its diameter unmeasurable. The exponential fit’s squared correlation coefficient (R²) and significance (P) are provided. **(B)** The disk expansion rate (cm/s) represents the change in disk diameter over time. **(C)** The perceived expansion rate (degrees/s) estimates the rate of change in the diameter of the disk over time, for observers placed either 10 cm (blue line), 20 cm (red line), 30 cm (yellow line) or 40 cm (purple line) away from the screen. The two horizontal dotted lines illustrate the range of expansion rates (35–350 degrees/s) previously determined to evoke locomotor responses [S1].

**Figure S4.**
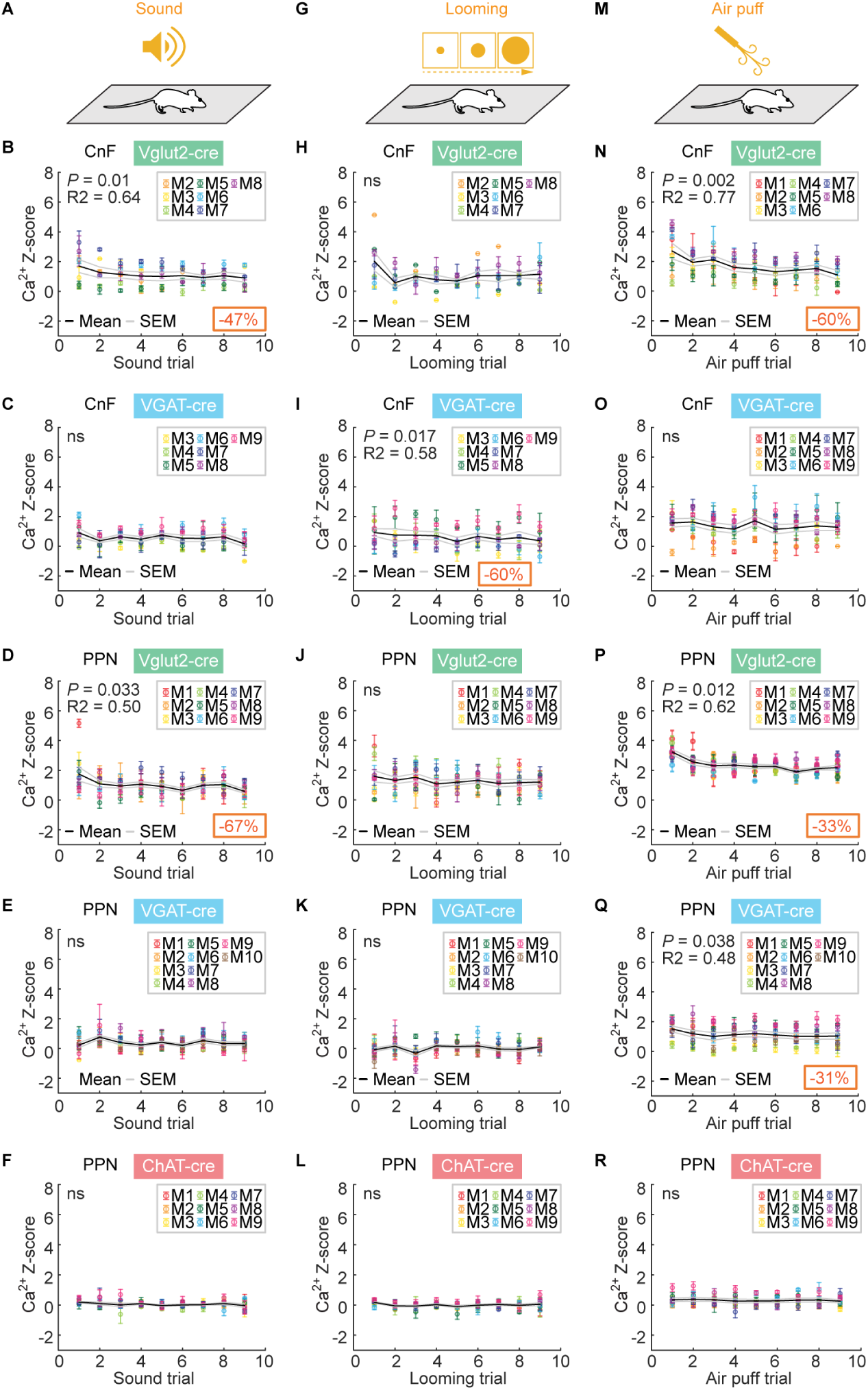
Habituation to sensory stimulations in CnF or PPN cells. **(A-R)** Mean calcium (Ca^2+^) responses recorded in the cuneiform nucleus (**CnF**) or pedunculopontine nucleus (**PPN**) of Vglut2-cre, VGAT-cre or ChAT-cre mice during the first 9 trials of 1-4 series of sound stimulations (A-F), visual looming stimulations (G-L), or air puff stimulations (M-R). For each trial number, the mean Ca^2+^ response ± SEM is illustrated. In each case, the existence of a linear relationship between trial number and Ca^2+^ signal amplitude was tested. When a fit is significant, the coefficient of correlation (R) and its significance (P) are illustrated, as well as the mean percentage decrease of the response from first to last trial (value in orange square) (ns, not significant, \**P* < 0.05, \*\**P* < 0.01, \*\*\**P* < 0.001, linear fit).

